# Neural Mechanisms of Intersensory Switching: Evidence for Delayed Sensory Processing and Increased Cognitive Effort

**DOI:** 10.1101/2024.11.26.625211

**Authors:** Theo Vanneau, Michael Quiquempoix, John J. Foxe, Sophie Molholm

## Abstract

Intersensory switching (IS), the ability to shift attention between different sensory systems, is essential for cognitive flexibility, yet leads to slower responses compared to repeating the same sensory modality. The underlying neural mechanisms of IS remain largely unknown. In this study, high-density EEG was used to investigate these mechanisms in healthy adults (n=53) performing a speeded reaction time (RT) task involving visual and auditory stimuli. Trials were categorized as Repeat (same preceding modality) or Switch (different preceding modality). Switch trials showed slower RTs and delayed sensory responses (N1 and P2 components). Furthermore, across both Repeat and Switch trials, RT correlated with the latency of these neural responses. Additionally, lower alpha-band inter-trial phase coherence (ITPC) in primary sensory regions was noted for Switch compared to Repeat trials, suggesting reduced efficiency of sensory processing. Greater induced theta activity over fronto-central scalp regions in Switch trials suggested increased cognitive control demands, potentially involving the anterior cingulate cortex (ACC). These findings reveal that IS is characterized by delayed sensory processing and heightened cognitive load, supporting a model where prior stimulus primes the sensory cortex for faster processing in Repeat trials, while Switch trials demand more cognitive resources for adjustment. The similarity of effects across both auditory and visual sensory modalities suggests that IS effects represent core features of sensory processing, potentially reflecting a fundamental, modality-independent mechanism of attentional switching across sensory domains.

## Introduction

Attentional capacity is limited, often requiring that attention be allocated selectively. The ability to switch attentional resources with changing task demands, or in response to novel sensory information, is fundamental to cognitive flexibility (Boulter, 1977; Klein, 1977; Lukas et al., 2010; Poole et al., 2021; Posner et al., 1976; Post & Chapman, 1991; Spence et al., 2001; Uppal et al., 2016; Wylie et al., 2003, 2004; Wylie et al., 2009). While numerous studies have shown a cost to switch between cognitive tasks, there is also evidence that simply changing sensory modality, even when the task itself does not change, likewise incurs a behavioral cost (see Tollner, 2009). For example, participants presented with stimulus pairs in which they were to ignore the first and perform a task on the second, responded slower when the S1-S2 sensory modalities differed (e.g. an auditory S1 followed by a visual S2) compared to when they matched (visual S1 followed by visual S2; (Turatto et al., 2002)). Intersensory switching costs (ISC) are also observed in paradigms in which mixed unisensory stimuli, all requiring the same simple motor response, are presented (Crosse et al., 2022; Shaw et al., 2020; Spence et al., 2001). In these cases, response time to a simple visual, auditory or tactile stimulus is longer when the previous stimulus was of a different sensory modality.

Despite the frequent occurrence of IS in everyday life, the neural mechanisms underlying ISC remain poorly understood. This stands in contrast to the well-studied task-switching costs, which have been linked to cortical regions involved in cognitive control including anterior cingulate cortex (ACC; (Johnston et al., 2007)), dorsolateral prefrontal cortex (DLPFC; (Sohn et al., 2000)), and inferior frontal gyrus (IFG; (Dove et al., 2000)). Task-switching costs are often attributed to increased cognitive demands, such as interpreting cues and reconfiguring stimulus-response mappings (e.g., when a participant is cued to respond to vowels instead of odd numbers in a letter-number task; (Monsell, 2003; Wylie & Allport, 2000; Wylie et al., 2003)). However, the behavioral cost of switching between sensory modalities, where there is no need to interpret a cue or change the cognitive or motor task, is not readily explained by increased cognitive demands.

Understanding the neural mechanisms underlying ISC is vital to shedding light on how the sensory systems operate and interact. At a theoretical level, two neural mechanisms may explain ISC. The first, *unisensory facilitation*, suggests that the preceding stimulus activates and biases the neural system towards that sensory modality, enhancing the response to subsequent stimuli of the same modality. This would not affect the processing of stimuli from different sensory modalities, so switch conditions are unaffected. This could be due, for example, to the stimulus evoked resetting of sensory specific attentional sampling (Plochl et al., 2022), to sensory priming, or predictive processes (Friston & Kiebel, 2009). The second, *intersensory competition*, assumes that competition among the sensory systems drives these behavioral patterns. This could be due to the involuntary reorienting of modality specific attention on a Switch trial (Bendixen et al., 2010), or alternatively, input into one sensory modality leading to inhibition of neural activity in other sensory modalities (Crabtree & Isaac, 2002; Cuppini et al., 2020). Of course, these processes of facilitation and competition are not mutually exclusive and could both be at play.

To date, there is a paucity of data that speaks to the neural mechanisms underlying ISCs. A few early studies focused on the P3, a cognitive component, with two such studies reporting larger P3 amplitudes after a modality change (Levit et al., 1973; Verleger & Cohen, 1978) and another reporting smaller P3s after a modality change (Rist & Cohen, 1987). Subsequent studies, focused on sensory processing, also yielded inconsistent findings. Gondan and colleagues (Gondan et al., 2007) found that Switch trials led to changes in the amplitude of sensory evoked responses, but in different directions depending on the sensory modality, with the auditory N1 response increasing and the visual N1 response decreasing. Tollner and colleagues (Tollner et al., 2009), on the other hand, found increased amplitude responses for switch compared to repeat trials for both tactile and visual stimuli over fronto-central scalp (but not parietal or lateral occipital scalp areas that are typically associated with somatosensory or visual sensory responses) in the 140-180ms timeframe. They attributed this to top-down activity in frontal cortical areas, due to attentional reweighting following switch trials. Thus, while ISCs are highly robust phenomena, the few studies directed at understanding the underlying neural processes have yielded somewhat inconsistent and inconclusive findings.

To address this significant gap in knowledge and better understand the interplay between the sensory and attentional systems, we analyzed high-density electrophysiological (EEG) data recorded from a large sample of adults engaged in a simple reaction-time task involving both visual and auditory stimuli under ‘Switch’ and ‘Repeat’ conditions. We performed both ERP and time-frequency analyses to test the involvement of sensory and top-down neural processing in IS, with an eye to addressing inconsistencies in the literature and to offer insights into the brain’s adaptive capacity for sensory processing across different modalities.

## Methods

### Participants and experimental procedure

The study was conducted with 55 healthy adults (20-63 yo, (26±7.39, Mean±Std); 25 males; note that except for 2 outliers, the age range was 20-33 (25.45±3.39)), all with normal hearing and vision (or corrected-to-normal). Participants sat 70 cm from a visual display (Dell UltraSharp 1704FPT) in an electrically shielded room (International Acoustics Company, Bronx, New York). The stimuli, controlled by Presentation software (Neurobehavioral Systems), included three types: a red disc (’Visual’), a 1000 Hz tone (’Audio’), and their simultaneous combination (’Audiovisual’). Participants were instructed to press a button as quickly as possible upon detecting any stimulus. The auditory stimulus was 1000Hz, 60ms tone presented binaurally (75 dB SPL). The visual stimulus was a red disc subtending to 1.5 degrees, displayed above a fixation cross. The audiovisual stimulus was a simultaneous presentation of both. Each trial presented a pseudo randomly chosen stimulus (A, V, or AV; represented equiprobably), with stimuli delivered through a speaker (JBL Duet) and displayed on a flat-panel LCD. A jittered interstimulus interval (1000-3000ms) reduced onset predictability. The task consisted of 1000 trials across 10 blocks. Button presses were recorded using a response pad (Logitech Wingman Precision Gamepad). Triggers indicating stimulus latency were sent from the PC acquisition computer via Presentation software.

All procedures were reviewed and approved by the Institutional Review Board at Einstein College of Medicine and conformed with the tenets for ethical conduct of human subjects’ research laid out in the Helsinki declaration. Participants provided written informed consent and were paid an hourly rate for their participation.

### EEG recordings & preprocessing

EEG data were recorded at a sampling rate of 512Hz using 160 channels BioSemi Active II system (using the CMS/DRL referencing system). Analyses were conducted in python (3.11) using MNE (originally for Minimum Norm Estimation)(Gramfort et al., 2013) and custom scripts available at https://github.com/tvanneau/Cross-sensory-switching. Bad channel detection was performed using the function NoisyChannels (with RANSAC) from the pyprep toolbox (Bigdely-Shamlo et al., 2015), if more than 15% of the channels were detected as bad, the participant was rejected (2 participants were rejected based on this criterion). Bad channels were interpolated using spline interpolation (Perrin et al., 1989). EEG was filtered using a FIR band-pass filter (0.5-40Hz), and Independent Component Analysis (ICA) on 1Hz high-pass EEG was used to identify and manually reject eye-related components (blinks/saccades). Epochs were created from −500 to +800ms around each stimulation with a baseline-correction from −200ms to stimulus onset. Epochs without a response from the participants were rejected. ‘Audiovisual’ trials were not used for the current analyses. For both visual and auditory stimulation, all the trials preceded by the same sensory modality were ‘Repeat’ trials and all the trials preceded by the other sensory modality were ‘Switch’ trials as illustrated in Figure 1 for visual stimuli and Figure 2 for auditory stimulation. For each sensory modality and for each individual, the 95th percentile of the RTs was calculated and trials associated with the 2.5% lower and upper bound responses rejected to avoid potential outliers. 5388 ‘Repeat’ visual trials (101/subject in average), 5302 ‘Switch’ visual trials (100/subject in average), 5370 ‘Repeat’ auditory trials (101/subject in average) and 5188 ‘Switch’ auditory trials (97/subject in average) were included for subsequent statistical analyses.

**Figure 1:**
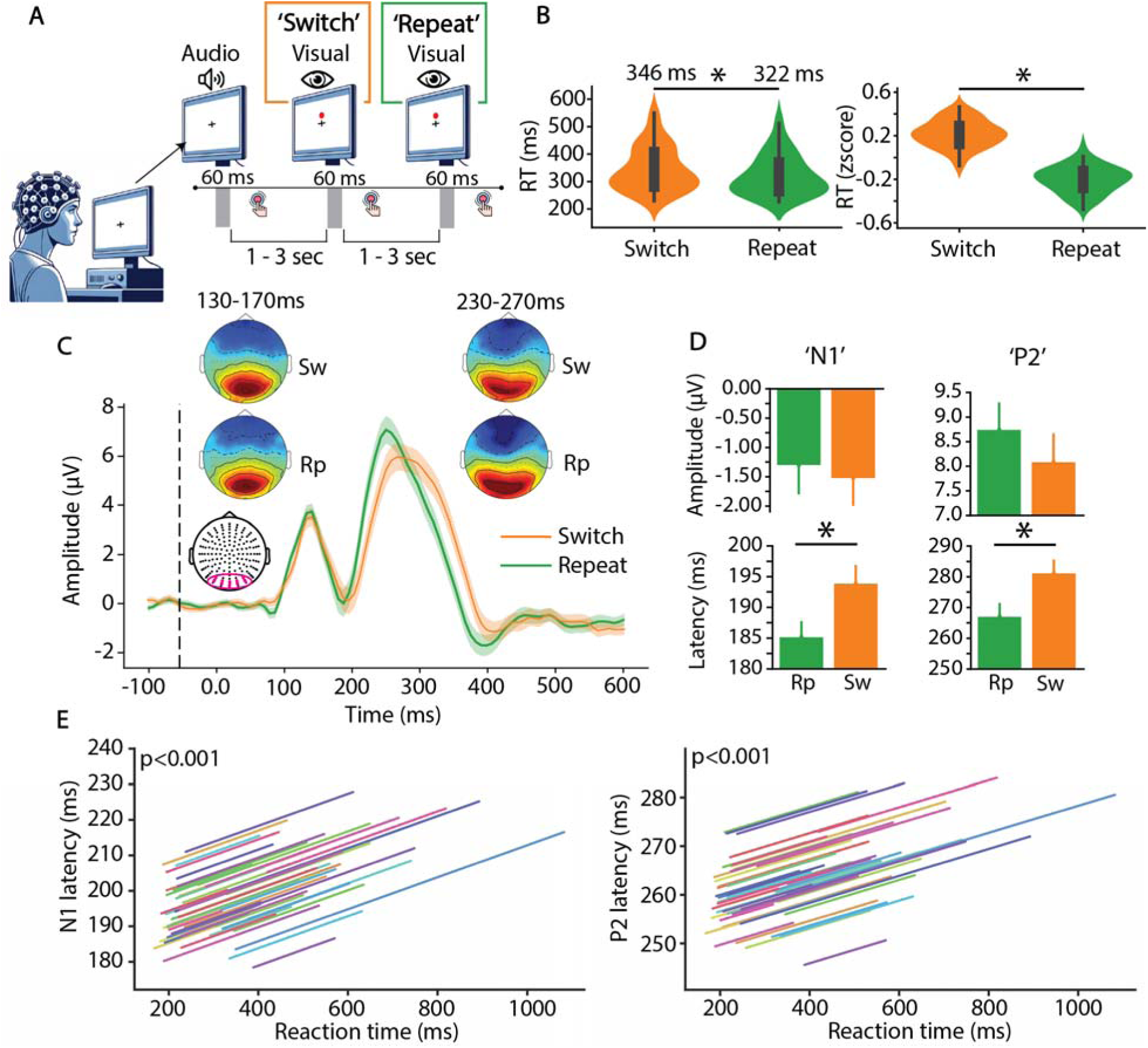
Increased N1 and P2 latency for visual-switch stimulations. **A**, Experimental paradigm illustrating a sequence of stimuli with a Switch and a Repeat trial for visual stimulation. **B**, on the left: average reaction time for Switch trials in orange and Repeat trials in green before normalization, the average RTs are displayed on the boxplot, on the right: after an individual z-score normalization. **C**, Topographical map of activity corresponding to the P1 and the P2 and ERP waveform averaged over a cluster of occipital electrodes for Switch (orange line) and Repeat trials (green line) with a frontal reference. **D**, Amplitude and latency of the N1 (minimum peak detection between 150 to 240ms) and for the P2 (maximum peak detection between 220 and 350ms) for Repeat (Rp, in green) and Switch trials (Sw, in orange). **E**, repeated-measure correlation (rmcorr) performed using single trials values between ERP parameters and RTs. Each line represents the data for an individual subject, where the slope of the lines is consistent across all subjects because the correlation coefficient is calculated collectively across all subjects. The central point of the line is located at the centroid of all the trials data for the corresponding subject. The length of each line reflects the combined variability of both variables (i.e., the Euclidean distance between data points for each trial). *Indicate significant differences assessed by LMMs (α=0.05).

**Figure 2:**
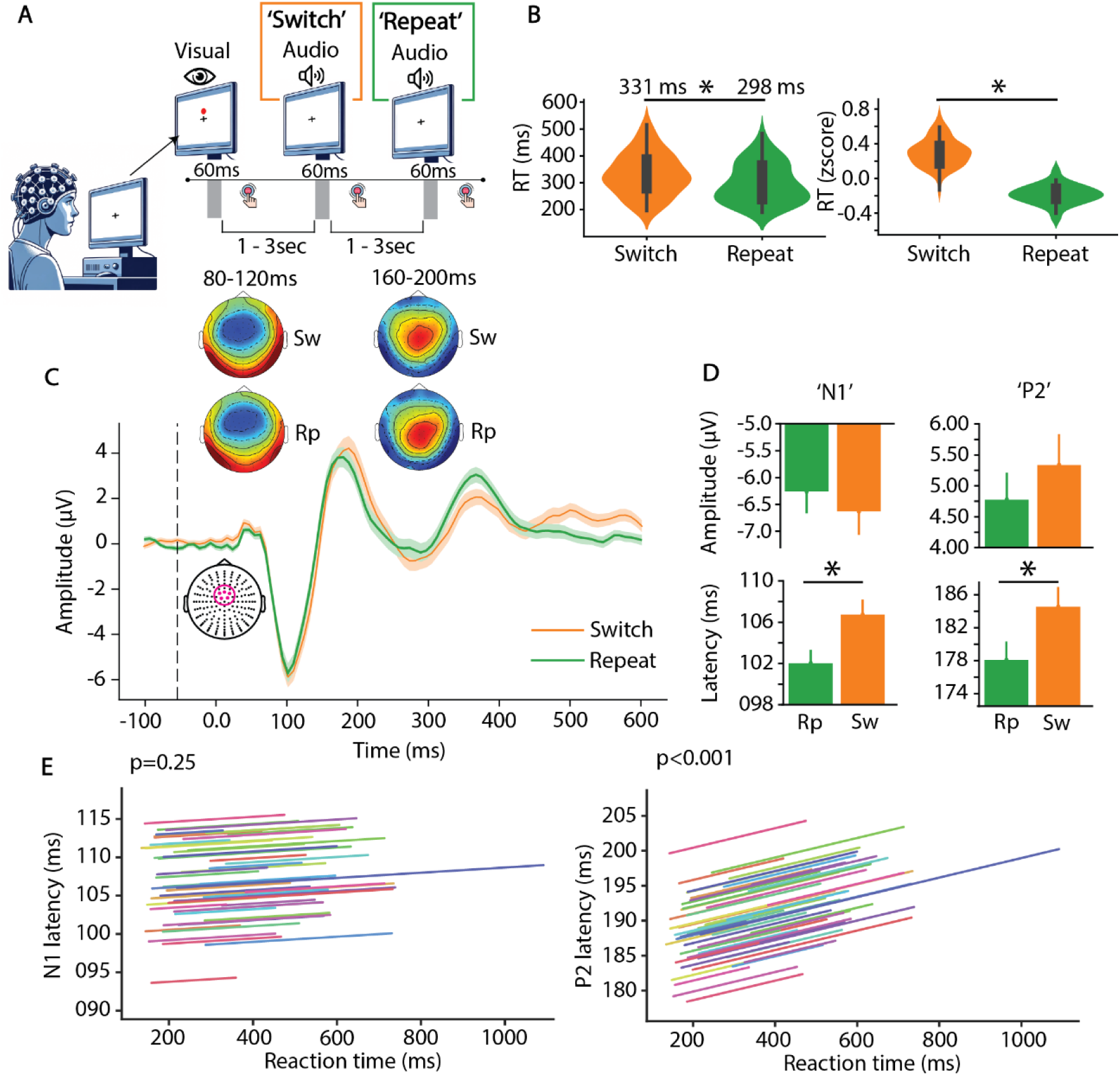
Increased N1 and P2 latency for auditory-switch stimulations. **A**, Experimental paradigm illustrating a sequence of stimuli with a Switch and a Repeat trial for auditory stimulation. **B**, on the left: average reaction time for Switch trials in orange and Repeat trials in green before normalization, the average RT are displayed on the boxplot, on the right: after an individual z-score normalization. **C**, Topographical map of activity corresponding to the N1 and the P2 and ERP for a cluster of frontal-central electrodes for Switch (orange line) and Repeat trials (green line) with a temporal reference. **D**, Amplitude and latency of the N1 (minimum peak detection between 70 to 150ms) and for the P2 (maximum peak detection between 150 and 240ms) for Repeat (Rp, in green) and Switch trials (Sw, in orange). **E**, repeated-measure correlation (rmcorr) performed using single trials values between ERP parameters and RTs. Each line represents the data for an individual subject, where the slope of the lines is consistent across all subjects because the correlation coefficient is calculated collectively across all subjects. The central point of the line is located at the centroid of all the trials data for the corresponding subject. The length of each line reflects the combined variability of both variables (i.e., the Euclidean distance between data points for each trial). *Indicate significant differences assessed by LMMs (α=0.05).

### ERP analysis

As can be seen from the scalp topographies and waveforms in Figures 1 & 2, both visual and auditory ERPs exhibited classical components as described in the literature (Foxe & Simpson, 2002; Picton et al., 1974). The visual response exhibited a dipolar activity focused on parieto-occipital channels and fronto-central channels, with a first positive peak around 100ms (the ‘P1’), a negative component around 190ms (the ‘N1’), and a second positive component around 250ms (the ‘P2’; Fig. 1). The auditory response showed dipolar activity that was focused over fronto-central and temporal scalp regions with a first positive component around 50ms (the P50), the N1 component around 100ms, and the P2 component around 180ms (Fig. 2). To maximize the visual sensory response for statistical analyses, visual ERPs were re-referenced to a fronto-central channel corresponding to FCz (whereas the ERP-related topographical maps on Figure 1 were plotted using common average reference). For calculation of amplitude and latency values for the visual ERP for Switch and Repeat trials we selected a cluster of 20 channels (from a 160-channel cap) over central and lateral parieto-occipital scalp where the VEP was greatest (see Fig. 1 and Supplementary Figure 1). Individual participant amplitude and latency for the visual N1 and P2 were extracted based on the minimum value between 150 to 240ms and the maximum value between 220 and 350ms respectively. To maximize the auditory sensory response for statistical analysis, auditory ERPs were re-referenced to a pair of symmetric temporo-parietal channels corresponding to TP7 and TP8 (the common average reference was used for the corresponding topographical maps displayed in Fig. 2). For calculation of amplitude and latency values for the auditory Switch and Repeat trials, data were extracted from a cluster of 5 channels over fronto-central scalp where the AEP was maximal (see Fig. 2 and Supplementary Figure 1). Individual participant amplitude and latency for the auditory N1 and P2 were extracted based on the minimum value between 70 and 150 and the maximum value between 150 and 240ms respectively. We also calculated difference waves by subtracting the average Repeat response from the average Switch response for each subject using the same cluster of channels as described above for the respective sensory modalities. To further assess potential differences in the neural generators of activity between the Switch and Repeat conditions and to pinpoint the timing of motor response identification, Current Source Density (CSD) maps were generated using the Laplacian second spatial derivative technique (Supplementary Figure 1, (Pascual-marqui et al., 1988)).

### Spectral analysis

In addition to ERP analysis, examining the spectral content of EEG signals provides valuable insights into the underlying neural mechanisms involved in cognitive processes. Spectral analysis involves decomposing the EEG signal into its constituent frequencies to study oscillatory activity that is not easily observable in time-domain data alone. Analyses were focused on the theta (3-7Hz) and alpha (7-13Hz) frequency bands. Oscillatory activity in the theta band is associated with cognitive control, working memory, and the coordination of neural networks (Cavanagh & Frank, 2014). The alpha band is related to attentional processes, sensory inhibition, and the regulation of cortical excitability (Foxe et al., 2014; Foxe & Snyder, 2011; Klimesch, 1999). While for ERP analysis specific references were used to increase the signal to noise ratio, for spectral analysis, a common-average reference was used to facilitate a direct comparison between visual and auditory induced oscillatory activity. Spectral analysis was performed using complex Morlet wavelet convolution (tfr_morlet function in MNE) between 2 and 40Hz with a FWHM in the spectral domain of 3.25Hz and 187ms in the temporal domain (Cohen, 2014) that correspond to: number of cycles = frequency / 2 (with frequency as a vector of frequency from 2 to 40 in steps of 0.19Hz). The non-phase locked power was calculated after subtracting the ERP from each trial for each individual (Kalcher & Pfurtscheller, 1995). For the total power and the non-phase locked power two different baseline normalization methods were used, that both gave the same results: A trial-by-trial baseline normalization and an average baseline normalization applied after averaging all the trials. Both methods were performed independently for every individual and using a −200ms to 0ms baseline period and a decibels (dB) normalization. The results presented in the figures are derived from the average baseline normalization.

### Statistical analysis

Statistical analyses were performed in Jamovi (v2.3.28) to compare Switch and Repeat trials for: Reaction times; ERP component amplitude and latency (Fig. 1 & 2); alpha-band ITPC peak amplitude and corresponding latency (Fig. 3); and fronto-central induced theta power peak amplitude and corresponding latency (Fig. 4). Analyses were conducted using linear mixed models (LMMs) to adjust for individual differences among subjects (with a degree of freedom of 1 for the fixed effects (Switch versus Repeat) and of 52 for the random effects (subjects) and with alpha set at 0.05), we reported for each test: the F-stats (‘F’, which indicate the significance of the fixed effect, here Switch versus Repeat), the Estimate (shows the size and direction of the effect, all tests were runed as ‘Switch-Repeat’) and the corresponding p-value. The normality of residuals was assessed using Kolmogorov-Smirnov and Shapiro-Wilk tests. To measure the correlation between ERP latencies and RTs while accounting for the within-subject effect, we used the repeated-measure correlation method (Bakdash & Marusich, 2017) on trial-by-trial data points with the rm_corr function of the pingouin toolbox in Python (Vallat, 2018). Repeated measures correlation is designed to estimate the strength and direction of the linear relationship between two variables while controlling for the variability due to differences between subjects. The function calculates the covariance between x and y within each subject. It then adjusts for the overall between-subject variability to isolate the within-subject correlation. The rm_corr function computes the correlation coefficient (r_rm) that reflects the relationship between x and y after accounting for the repeated measures design.

**Figure 3:**
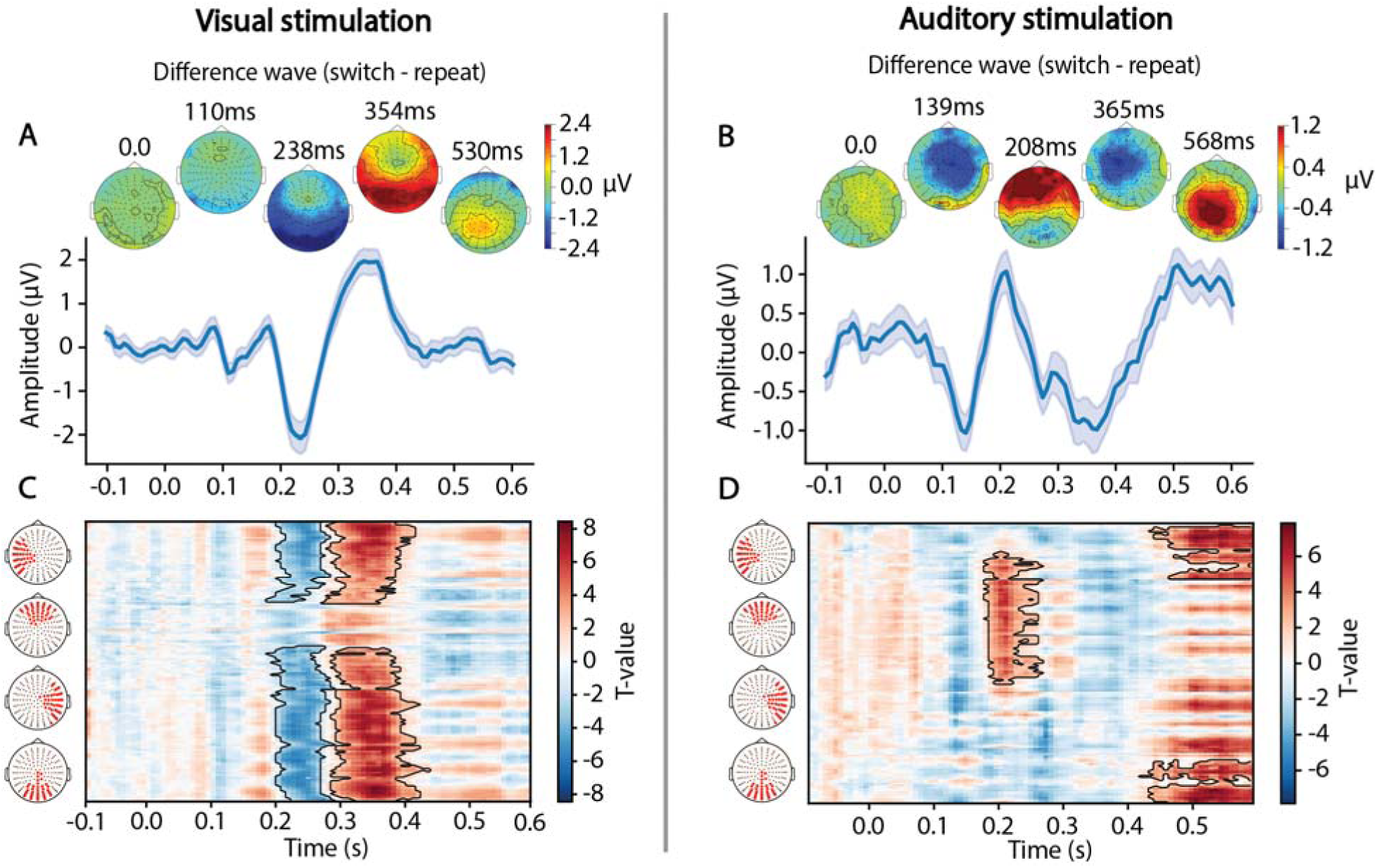
Spatio-temporal analysis of difference waves in visual and auditory switch trials: evidence of delayed N1 and P2 latencies. **A**, Difference wave between visual responses in switch and repeat trials (same cluster of channels as in Fig. 1), accompanied by the corresponding topographical map highlighting regions of interest and the timing of key components. **B**, Difference wave between auditory responses in switch and repeat trials (same cluster of channels as in Fig. 2), with the associated topographical map indicating relevant regions and timings. **C**, Spatio-temporal cluster-based permutation test results for the visual difference wave (see Methods), with significant clusters over time and across channels outlined in a black trace. The t-values for each channel and time point are displayed in the plot. **D**, Spatio-temporal cluster-based permutation test results for the auditory difference wave, as described for C.

**Figure 4:**
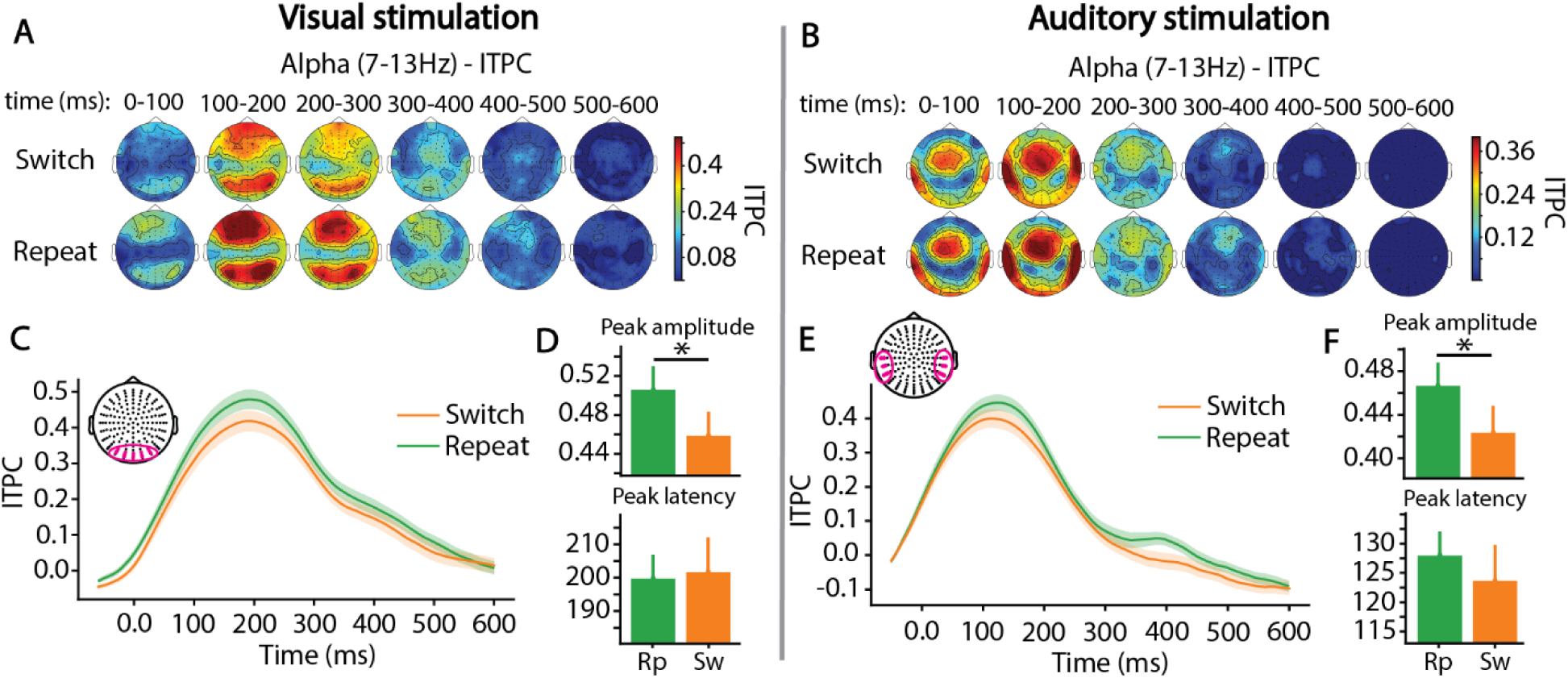
Higher alpha-band ITPC in response to repeat trials. **A**, Topographical map of the ITPC in the alpha-band (7-13Hz) in response to visual stimulation for Switch (upper row) and Repeat (bottom row) trials from 0 to 600ms. **B,** same as **A** but in response to auditory stimulation. **C**, time course of the alpha-band ITPC for a cluster of occipital electrodes for Switch (orange line) and Repeat (green line) trials. **D**, corresponding maximum peak-amplitude and associated latency extracted for each individual for Repeat (Rp, green) and Switch (Sw, orange) trials. **E** and **F**, same as C and D for a cluster of electrodes over bilateral temporal scalp regions in response to auditory stimulation. * Indicate significant differences assessed by LMMs (α=0.05).

In a further exploratory analysis phase, and in order to take full advantage of the richness of the dataset herein, spatio-temporal cluster-based permutation tests (Maris & Oostenveld, 2007) were performed over the full set of channels and time-points. The function permutation_cluster_1samp_test was used to compare the Switch minus Repeat difference wave to 0 (no difference between conditions). We used the statistic from one-sample t-test with 1000 permutations and a threshold corresponding to α=0.05. The idea is that the conditions are randomly shuffled (permuted) many times (e.g., 1000 permutations). For each permutation, the same steps (statistical test and cluster formation) are performed. For each permutation, the largest cluster-level statistic is recorded, creating a distribution of cluster-level statistics under the null hypothesis (no difference between conditions). The cluster-level statistics from the original data (before permutation) are compared to the distribution of cluster-level statistics generated from the permutations. The p-value for each cluster is determined based on where it falls in the permutation distribution. Clusters with p-values below a certain threshold (e.g., 0.05) are considered statistically significant. The permutation test inherently controls multiple comparisons by focusing on clusters rather than individual points. This means that the method reduces the risk of false positives due to the large number of tests conducted across time points and channels.

## Results

### Common delayed N1 and P2 components for visual and auditory Switch trials

The first part of this study focuses on behavioral and ERP analysis. We first consider ISCs for visual stimuli. In this context, Switch trials were visual stimulations preceded by auditory stimulation, whereas Repeat trials were preceded by visual stimulation (Fig. 1A). Reaction times (RTs) to Switch trials were significantly slower compared to Repeat trials (Fig. 1B; F=93.4, Estimate=24.0 and p<0.001), even after individual z-score normalization to account for inter-individual variability in RT (F=295.0, Estimate=0.41 and p<0.001).

The ERP-based scalp topographies for Visual Repeat and Switch trials were similar between the two conditions (Fig. 1C; see Methods for a description of the components), even when using the Current Source Density (CSD) signal to improve localization (Supplementary Figure 1). Statistical analyses revealed a significant difference in N1 and P2 latencies, which reflected earlier peak responses for Repeat compared to Switch trials (Fig. 1D; for the N1: F= 12.8, Estimate= 8.70 and p<0.001; for the P2: F= 11.9, Estimate= 14.2 and p= 0.001). However, no significant differences were found in N1 amplitude (F= 0.59, Estimate = −0.22, p = 0.49) or P2 amplitude (F= 7.85, Estimate= −0.66, p= 0.09) between Switch and Repeat trials.

### Correlation between visual evoked brain responses and reaction times

The four parameters of interest for each trial for each participant were extracted and used a repeated-measures correlation method, a statistical technique that measures the association between two variables while accounting for the within-subject design (see Methods). This revealed a small but significant positive correlation between RTs and both N1 (Fig. 1E; r= 0.11, df= 10636, p<0.001, power= 1.0) and P2 latency (r= 0.07, df= 10636, p<0.001, power= 1.0), and a negative correlation between RTs and both N1 (Supplementary Figure 2; r= −0.06, df= 10636, p<0.001, power= 0.99) and P2 amplitude (r= −0.12, df= 10636, p<0.001, power= 1.0).

We next considered ISC for auditory stimuli. In this context, Switch trials were now preceded by visual stimulation whereas Repeats were preceded by another auditory stimulation (Fig. 2A). As with the visual stimulation, RTs associated with Switch trials were slower both for raw values (Fig. 2B; F= 127.0, Estimate= 33.5 and p<0.001) and after individual z-score normalization (F= 315.0, Estimate= 0.48 and p<0.001).

The ERP-related scalp topographies for Auditory Repeat and Switch trials were similar across both conditions (Fig. 1C; see Methods for a description of the components), even when utilizing CSD signal to enhance localization (Supplementary Figure 1). Statistical analyses revealed a significantly delayed latency for Switch compared to Repeat trials for both the N1 (Fig. 2D; F= 8.76, Estimate= 4.72 and p= 0.005) and the P2 (F= 11.3, Estimate= 6.49 and p= 0.001) components. However, no significant differences were observed in the amplitudes of the N1 (Fig. 2E; F= 2.74, Estimate= −0.37, p= 0.23) or the P2 (F= 6.35, Estimate= −0.56, p= 0.12) between conditions.

### Correlation between auditory evoked brain responses and reaction times

Correlation analysis failed to reveal a significant relationship between RT and either N1 amplitude or latency (Supplementary Figure 3 for amplitude: r= 0.005, df= 10504, p= 0.6, power= 0.08; Fig. 2E for latency: r= 0.01, df= 10504, p= 0.25 and power= 0.2). However, the repeated-measure correlation was significant between RTs and both the latency (r= 0.05, df= 10504, p<0.001, power= 0.99) and amplitude (r= 0.06, df= 10504, p<0.001, power= 0.99) of the P2, which corresponds to the latency of the visual N1.

To further explore differences between the Switch and Repeat conditions, spatio-temporal cluster-based permutation tests were performed on the average difference wave between Switch and Repeat trials, computed for each subject for each of the sensory modalities. For visual responses, the difference wave displayed a negative deflection peaking at 238ms, followed by a positive deflection peaking at 358ms. The corresponding topographical pattern indicated that these differences originated primarily from regions over posterior scalp associated with visual sensory processing, particularly the occipital channels (Fig. 3A). Cluster-based permutation statistics revealed that the negative values observed between 200 and 300ms were significantly different across occipital and temporal channels (Fig. 3B). Notably, these negative values, which are shifted relative to the average N1 peak (around 190ms), support the conclusion that the N1 latency is delayed in Switch trials. Similarly, the analysis identified significant positive values occurring after the P2 peak latency, between 300 and 400ms. For auditory stimulation, the difference wave exhibited a negative deflection peaking at 139ms, followed by a positive deflection at 208ms, and another negative going deflection at 365ms. As with visual stimulation, the topographical pattern of this difference wave suggested that the observed differences primarily originated from scalp regions associated with auditory processing, particularly over frontal-central channels, likely influenced by the temporal reference (see Methods). The statistical analysis revealed significant positive values between 200 and 280ms across frontal-central channels, which are shifted relative to the average P2 peak (around 180ms), lending further support to an interpretation of delayed latency of the P2 for Switch trials. In contrast, the negative values associated with the delayed N1 (around 130ms on the cluster plot) were not statistically significant.

### Higher alpha-band inter-trial phase coherence for Repeat trials

Inter-trial phase coherence (ITPC) reflects the consistency of the phase of neural oscillations across trials. This consistency can reflect enhanced precision of sensory processing, which is crucial for accurate perception and response. While we analyzed the ITPC for the theta (3-7Hz), alpha (7-13Hz) and beta (13-30Hz) frequency bands, only the alpha-band ITPC revealed consistent differences between Switch and Repeat trials. The topographical map of the alpha-band ITPC highlights higher values for each respective primary sensory region for each type of stimulation, occipital scalp for visual stimulation (Fig. 4A), and temporal scalp for auditory stimulation (Fig. 4B). The ITPC increased relative to baseline in response to visual stimulation over occipital scalp, with a peak around 200ms for both Repeat and Switch trials (Fig. 4C). Whereas the latency of this peak did not differ between conditions (Fig. 4D; F= 0.04, Estimate= 1.92 and p= 0.84), alpha-ITPC was significantly higher for Repeat compared to Switch trials (F= 5.55, Estimate= −0.04 and p= 0.02). The same pattern was observed over temporal scalp regions in response to auditory stimulation with an alpha-ITPC peak around 125ms (Fig. 4E) that did not significantly differ in terms of latency between conditions (Fig. 4F; F= 0.143, Estimate= 0.003 and p= 0.70), but as for visual stimulation, ITPC peak-amplitude was significantly higher for Repeat trials (F= 4.69, Estimate= −0.05 and p= 0.03).

### Delayed and increased theta activity over frontal cortex for Switch trials

Theta activity in frontal regions, such as the anterior cingulate cortex (ACC) and prefrontal cortex (PFC), is often associated with top-down control mechanisms (Min et al., 2010; Helfrich et al., 2019). This includes the modulation of sensory processing by cognitive factors such as attention, expectation, and task demands. To determine potential involvement of frontal theta activity, we analyzed the non-phase locked power (‘induced oscillations’, obtained after subtraction of the stationary ERP to each trial) in the theta band (3-7Hz) in response to Switch and Repeat conditions for both visual and auditory stimulation. By focusing on induced oscillations rather than total power, we ensured that the observed differences in power did not merely reflect differences in stimulus evoked ERPs. For both sensory modalities and for both Switch and Repeat trials, induced theta power over frontal-central scalp showed a remarkably similar spatio-temporal pattern (Fig. 5A, B) with a maximum around 300ms (Fig. 5C and 5E). Statistical analysis revealed that the amplitude and peak latency of this activity was greater for Switch compared to Repeat trials for both visual (Fig. 5D; Peak amplitude: F= 11.3, Estimate= 0.2 and p= 0.001; Peak latency: F= 4.93, Estimate= 17.7 and p= 0.03) and auditory conditions (Fig. 5F; Peak amplitude: F= 19.4, Estimate= 0.04 and p<0.001; Peak latency: F= 6.63, Estimate= 28.9 and p= 0.01).

**Figure 5:**
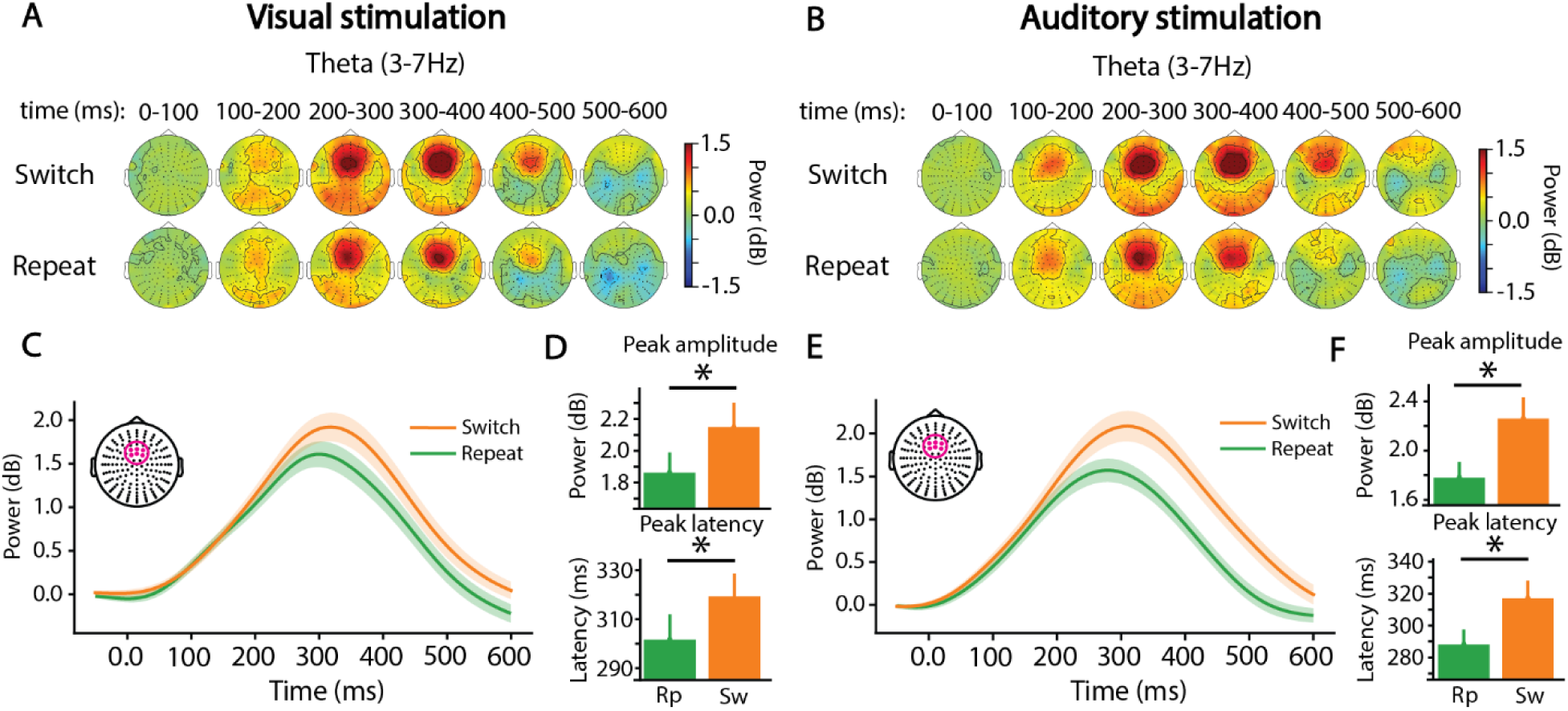
Higher activation of theta activity over centro-frontal scalp to switch trials. **A**, Topographical map of the induced theta-band power in response to Switch (upper row) and Repeat (bottom row) visual stimulation and **B**, for auditory stimulation. **C**, time course of the theta-band induced power for a cluster of frontal-central electrodes for Switch (orange line) and Repeat (green line) visual stimulation. **D**, corresponding maximum peak-amplitude and associated latency extracted for each individual for Repeat (Rp, green) and Switch (Sw, orange) trials. **E** and **F**, same as **C** and **D** for the same fronto-central cluster of electrodes but in response to Switch and Repeat auditory stimulation. * Indicate significant differences assessed by LMMs (α=0.05).

## Discussion

Intersensory switching (IS) is crucial for flexibly interacting with the environment, but often leads to delayed responses compared to repeat trials within the same sensory modality (Boulter, 1977; Crosse et al., 2022; Klein, 1977; Lukas et al., 2010; Poole et al., 2021; Posner et al., 1976; Post & Chapman, 1991; Shaw et al., 2020; Spence et al., 2001). Although IS is a common occurrence in daily life, the neural mechanisms behind the increased reaction times (RTs) associated with switching between modalities are not well understood. In this study, we used visual and auditory stimuli in conjunction with high-density EEG to explore the neural consequences of IS in a large cohort of healthy adults (n=53).

Findings reveal that not only do RTs increase during modality switches, but there also appears to be a common neural pattern across sensory modalities that accounts for this delay. This pattern is characterized by both increased sensory processing latency, reflected in delayed ERP components, and greater cognitive effort, indicated by heightened fronto-central theta activity, suggesting similar brain mechanisms underlying IS for both visual and auditory modalities. Furthermore, for both sensory modalities, there was a significant correlation between ERP latency and reaction time, reinforcing the thesis that ISC partly stems from relatively delayed sensory processing on Switch trials (Fellinger et al., 2011; Klimesch et al., 2007). Interestingly, this correlation was particularly strong for similar brain response latencies (180-220 ms, corresponding to the visual N1 and auditory P2) across both sensory modalities, indicating a potentially important temporal relationship between sensory processing and motor responses. Further evidence of altered sensory processing for Switch and Repeat trials is to be found in the relative decrease in peak alpha-band Inter-Trial Phase Coherence (ITPC) for Switch trials. Alpha-phase dynamics are thought to regulate cortical excitability, thereby influencing sensory processing (Klimesch et al., 2007). Higher ITPC in the alpha band for Repeat trials could indicate that the brain more effectively anticipates the upcoming stimulus, leading to a more synchronized and efficient neural response when the same sensory modality is presented consecutively, as opposed to when there is a switch. According to this explanation, when switching between modalities, the lack of such a priming effect leads to less efficient sensory processing and increased cognitive effort, which is consistent with the finding of heightened theta band activity on switch trials. The anterior cingulate cortex (ACC), known for its role in cognitive control, likely contributes to this process (Botvinick et al., 2004; Carter et al., 1999; Carter et al., 1998; Holroyd & Coles, 2002; Orr & Weissman, 2009; Shenhav et al., 2013), possibly reflecting the effort required in switching attentional resources to the alternate modality. We hypothesize that on repeat trials the preceding sensory modality primes sensory networks, leading to more efficient sensory processing and reduced cognitive effort. When a different sensory modality is introduced, it results in a delayed sensory response and ACC activation, reflecting increased cognitive effort and the potential reconfiguration of attention and updating predictions toward the new sensory modality.

It is interesting to note that the increased frontal theta activity supports the dimension-weighting hypothesis proposed by Töllner et al. (2009), suggesting that intersensory switching (IS) may work similarly to dimension switching within a single sensory modality. According to this hypothesis, when the task-relevant dimension (e.g., color or orientation) changes across trials, reaction times are slowed, just as they do when switching between sensory modalities (Dyson & Quinlan, 2002; Found & Muller, 1996; Muller et al., 2003). Repeating the same dimension leads to faster detection, while switching features requires a reallocation of attentional resources, resulting in slower responses. At the core of this idea is the dynamic allocation of attention based on which feature or dimension is relevant to the task (Liesefeld & Muller, 2019). In dimension switching, attention shifts from previously important features to new ones as the task changes. Research using ERPs and fMRI has shown that this reallocation process involves the anterior cingulate cortex, which contributes to these attentional shifts (Liston et al., 2006; Luks et al., 2002). In our study, the increased frontal theta activity during modality switching may similarly reflect a shift in attention between different sensory modalities.

A critical question that remains is whether the neural mechanisms underlying ISCs reflect a facilitatory effect of repeating the modality or an interference effect when switching modalities. From a practical standpoint, the nature of ERP evaluation and the relative comparison between Switch and Repeat trials make it difficult to disentangle these effects. The differences in ERP component latencies could result from either the facilitatory effect of repeating the target modality or an interference mechanism during switching. This issue arises because there is no standalone ‘reference’ brain response for each modality, as ERP calculations rely on averaging responses across multiple trials, inherently introducing repetition and conflating the effects of repetition with those of switching. One approach to disentangle facilitation versus interference could be to present longer sequences of repeat trials. If the effect is primarily due to facilitatory processes driven by expectancy, the latency of ERP components and corresponding reaction-times might decrease with successive repetitions, reaching a ceiling effect after a certain number of stimuli. Conversely, if the effect is driven by unexpected stimuli, the increase in latency for switch trials should become more pronounced as the number of preceding repetitions within the same sensory modality increases. However, in studies using longer sequences of repeated stimuli and considering the corresponding physiology, it will be important to also consider adaptation mechanisms that could reduce the amplitude of sensory responses for longer sequences of repeating modality (Andrade et al., 2015; Liston et al., 2006; Luks et al., 2002; Uppal et al., 2016).

The results presented here are consistent with previous behavioral research from our group, which demonstrated that much of the multisensory redundant signal effect (RSE) — the well-known multisensory speeding effect shown by violation of Millers race model (Megevand et al., 2013; Miller, 1982; Molholm et al., 2002)— can be explained by IS when comparing reaction times to multisensory versus unisensory stimuli (Crosse et al., 2022; Shaw et al., 2020). The typical multisensory RSE paradigm involves random sequencing of unisensory and bisensory inputs in a mixed design, which introduces an alternative explanation based on IS effects. In this design, participants frequently switch between sensory modalities (e.g., from responding to visual to auditory stimuli), leading to slower reaction times for unisensory trials. As a result, the apparent faster response to multisensory stimuli is not due to true speeding but rather to the slower responses on unisensory trials, creating a false RSE. This issue can also extend to neural findings associated with multisensory integration (Molholm et al., 2002). If a study combines unisensory switch and repeat trials to compare with multisensory responses, the results may be biased (Gondan et al., 2007) because the unisensory trials include switching costs, which may be absent in multisensory trials.

## Conclusion

Our study investigated the neural mechanisms underlying ISCs using visual and auditory stimuli in healthy adults. We found that RTs were shorter, ERP latencies (N1 and P2) were decreased, and alpha-band ITPC was higher for repeat trials compared to switch trials, indicating more efficient sensory processing during repeat trials. The increased fronto-central theta activity observed during switch trials suggests greater cognitive effort when switching between sensory modalities, involving the ACC. These results support a priming-expectancy hypothesis, where prior exposure to a modality biases the sensory network towards that same modality, facilitating more efficient processing on repeat trials. In contrast, switching modalities leads to delayed processing and heightened ACC activity, reflecting increased cognitive effort. Additionally, our findings align with previous research, which shows that multisensory integration effects, such as the RSE, are influenced by IS.

## Data & Code availability

The raw EEG data that supports the findings of this study are available from the corresponding author upon reasonable request. Analyses were conducted using custom Python script available on GitHub (https://github.com/tvanneau/Cross-sensory-switching).

## Author contributions

S.M and J.J.F conceived the study. T.V preprocessed the data and analyzed the data under the supervision of S.M and with input from MQ. T.V wrote the first draft of the manuscript. M.Q, J.J.F and S.M edited the manuscript.

---

**Supplementary figure 1.**
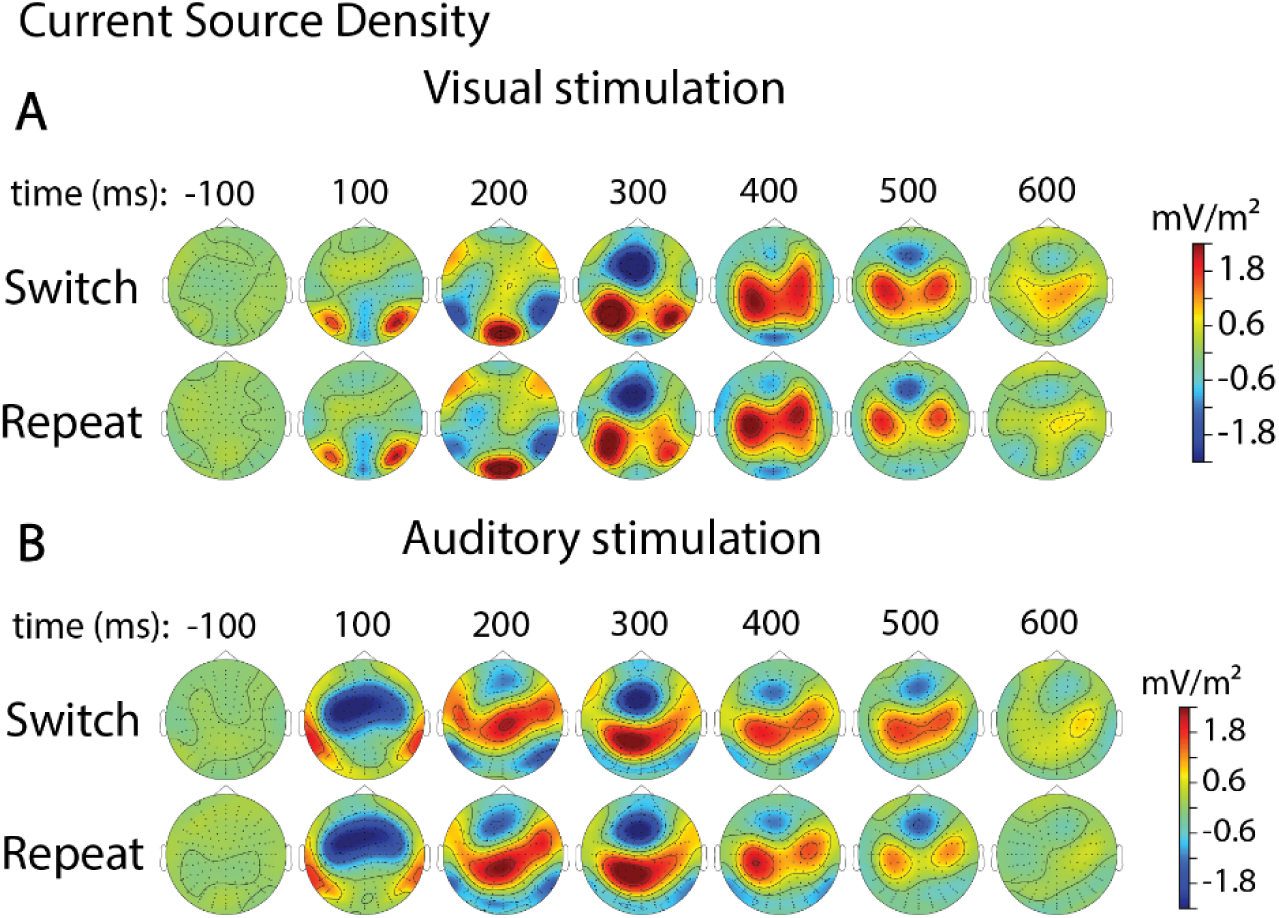
Current Source Density of visual and auditory evoked potential. Current Source Density of the event-related potential for Switch (upper line) and Repeat trials (bottom line) from −100 to 600ms with 100ms steps.

**Supplementary figure 2.**
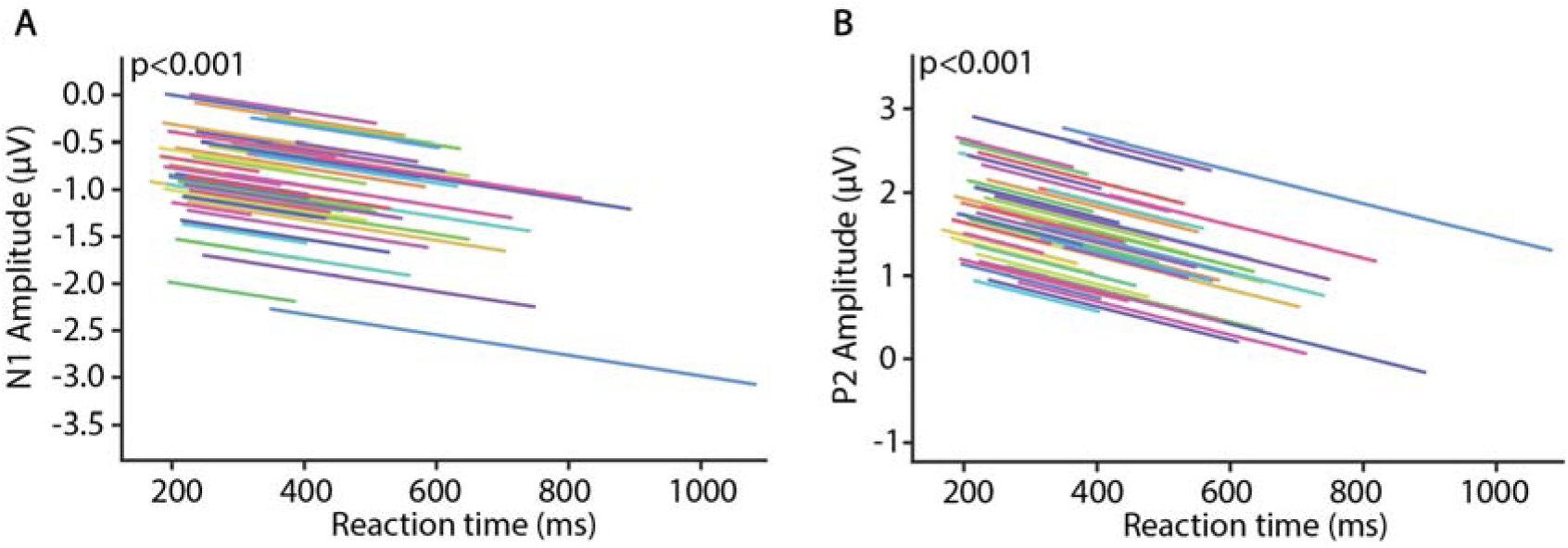
Repeated-measure correlation between visual-ERP amplitude and reaction time. Repeated-measure correlation (rmcorr) performed using single trials values between ERP parameters and RTs. Each line represents the data for an individual subject, where the slope of the lines is consistent across all subjects because the correlation coefficient is calculated collectively across all subjects. The central point of the line is located at the centroid of all the trials data for the corresponding subject. The length of each line reflects the combined variability of both variables (i.e., the Euclidean distance between data points for each trial).

**Supplementary figure 3.**
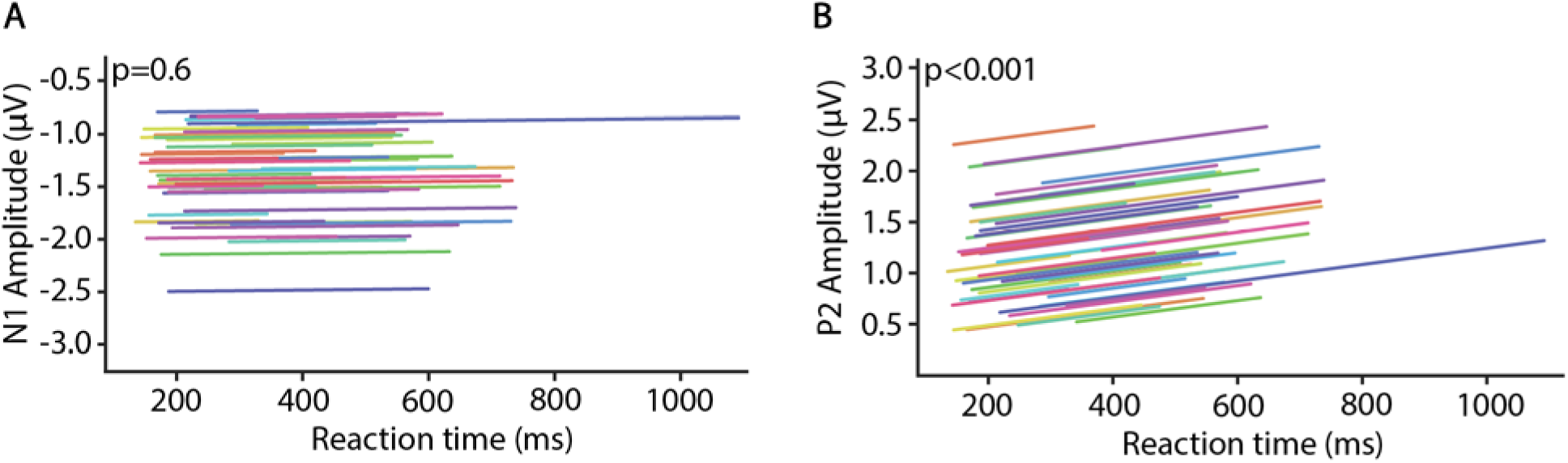
Repeated-measure correlation between auditory-ERP amplitude and reaction time. Repeated-measure correlation (rmcorr) performed using single trials values between ERP parameters and RTs. Each line represents the data for an individual subject, where the slope of the lines is consistent across all subjects because the correlation coefficient is calculated collectively across all subjects. The central point of the line is located at the centroid of all the trials data for the corresponding subject. The length of each line reflects the combined variability of both variables (i.e., the Euclidean distance between data points for each trial).

## References

Andrade, G. N., Butler, J. S., Mercier, M. R., Molholm, S., & Foxe, J. J. (2015). Spatio-temporal dynamics of adaptation in the human visual system: a high-density electrical mapping study. Eur J Neurosci, 41(7), 925–939. 10.1111/ejn.12849

Bakdash, J. Z., & Marusich, L. R. (2017). Repeated Measures Correlation. Front Psychol, 8, 456. 10.3389/fpsyg.2017.00456

Bendixen, A., Grimm, S., Deouell, L. Y., Wetzel, N., Madebach, A., & Schroger, E. (2010). The time-course of auditory and visual distraction effects in a new crossmodal paradigm. Neuropsychologia, 48(7), 2130–2139. 10.1016/j.neuropsychologia.2010.04.004

Bigdely-Shamlo, N., Mullen, T., Kothe, C., Su, K. M., & Robbins, K. A. (2015). The PREP pipeline: Standardized preprocessing for large-scale EEG analysis. Frontiers in Neuroinformatics, 9(JUNE), 1–19. 10.3389/fninf.2015.00016

Botvinick, M. M., Cohen, J. D., & Carter, C. S. (2004). Conflict monitoring and anterior cingulate cortex: an update. Trends Cogn Sci, 8(12), 539–546. 10.1016/j.tics.2004.10.003

Boulter, L. R. (1977). Attention and reaction times to signals of uncertain modality. J Exp Psychol Hum Percept Perform, 3(3), 379–388. 10.1037//0096-1523.3.3.379

Carter, C. S., Botvinick, M. M., & Cohen, J. D. (1999). The Contribution of the Anterior Cingulate Cortex to Executive Processes in Cognition. Rev Neurosci, 10(1), 49–58. doi:10.1515/REVNEURO.1999.10.1.49

Carter, C. S., Braver, T. S., Barch, D. M., Botvinick, M. M., Noll, D., & Cohen, J. D. (1998). Anterior cingulate cortex, error detection, and the online monitoring of performance. Science, 280(5364), 747–749. 10.1126/science.280.5364.747

Cavanagh, J. F., & Frank, M. J. (2014). Frontal theta as a mechanism for cognitive control. Trends Cogn Sci, 18(8), 414–421. 10.1016/j.tics.2014.04.012

Cohen, M. X. (2014). Fluctuations in oscillation frequency control spike timing and coordinate neural networks. Journal of Neuroscience, 34(27), 8988–8998. 10.1523/JNEUROSCI.0261-14.2014

Crabtree, J. W., & Isaac, J. T. (2002). New intrathalamic pathways allowing modality-related and cross-modality switching in the dorsal thalamus. Journal of Neuroscience, 22(19), 8754–8761. 10.1523/JNEUROSCI.22-19-08754.2002

Crosse, M. J., Foxe, J. J., Tarrit, K., Freedman, E. G., & Molholm, S. (2022). Resolution of impaired multisensory processing in autism and the cost of switching sensory modality. Communications Biology, 5(1). 10.1038/s42003-022-03519-1

Cuppini, C., Ursino, M., Magosso, E., Crosse, M. J., Foxe, J. J., & Molholm, S. (2020). Cross-sensory inhibition or unisensory facilitation: A potential neural architecture of modality switch effects. Journal of Mathematical Psychology, 99. 10.1016/j.jmp.2020.102438

Dove, A., Pollmann, S., Schubert, T., Wiggins, C. J., & von Cramon, D. Y. (2000). Prefrontal cortex activation in task switching: an event-related fMRI study. Brain Res Cogn Brain Res, 9(1), 103–109. 10.1016/s0926-6410(99)00029-4

Dyson, B. J., & Quinlan, P. T. (2002). Within- and between-dimensional processing in the auditory modality. Journal of Experimental Psychology: Human Perception and Performance, 28(6), 1483–1498. 10.1037/0096-1523.28.6.1483

Fellinger, R., Klimesch, W., Gruber, W., Freunberger, R., & Doppelmayr, M. (2011). Pre-stimulus alpha phase-alignment predicts P1-amplitude. Brain Res Bull, 85(6), 417–423. 10.1016/j.brainresbull.2011.03.025

Found, A., & Muller, H. J. (1996). Searching for unknown feature targets on more than one dimension: investigating a “dimension-weighting” account. Percept Psychophys, 58(1), 88–101. 10.3758/bf03205479

Foxe, J. J., Murphy, J. W., & De Sanctis, P. (2014). Throwing out the rules: Anticipatory alpha-band oscillatory attention mechanisms during task-set reconfigurations. Eur J Neurosci, 39(11), 1960–1972. 10.1111/ejn.12577

Foxe, J. J., & Simpson, G. V. (2002). Flow of activation from V1 to frontal cortex in humans. A framework for defining “early” visual processing. Exp Brain Res, 142(1), 139–150. 10.1007/s00221-001-0906-7

Foxe, J. J., & Snyder, A. C. (2011). The role of alpha-band brain oscillations as a sensory suppression mechanism during selective attention. Frontiers in Psychology, 2(JUL). 10.3389/fpsyg.2011.00154

Friston, K., & Kiebel, S. (2009). Predictive coding under the free-energy principle. Philos Trans R Soc Lond B Biol Sci, 364(1521), 1211–1221. 10.1098/rstb.2008.0300

Gondan, M., Vorberg, D., & Greenlee, M. W. (2007). Modality shift effects mimic multisensory interactions: an event-related potential study. Exp Brain Res, 182(2), 199–214. 10.1007/s00221-007-0982-4

Gramfort, A., Luessi, M., Larson, E., Engemann, D. A., Strohmeier, D., Brodbeck, C., Goj, R., Jas, M., Brooks, T., Parkkonen, L., & Hamalainen, M. (2013). MEG and EEG data analysis with MNE-Python. Front Neurosci, 7, 267. 10.3389/fnins.2013.00267

Holroyd, C. B., & Coles, M. G. H. (2002). The neural basis of human error processing: Reinforcement learning, dopamine, and the error-related negativity. Psychological Review, 109(4), 679–709. 10.1037//0033-295x.109.4.679

Johnston, K., Levin, H. M., Koval, M. J., & Everling, S. (2007). Top-down control-signal dynamics in anterior cingulate and prefrontal cortex neurons following task switching. Neuron, 53(3), 453–462. 10.1016/j.neuron.2006.12.023

Kalcher, J., & Pfurtscheller, G. (1995). Discrimination between phase-locked and non-phase-locked event-related EEG activity. Electroencephalogr Clin Neurophysiol, 94(5), 381–384. 10.1016/0013-4694(95)00040-6

Klein, R. M. (1977). Attention and visual dominance: a chronometric analysis. J Exp Psychol Hum Percept Perform, 3(3), 365–378. 10.1037//0096-1523.3.3.365

Klimesch, W. (1999). EEG alpha and theta oscillations reflect cognitive and memory performance: a review and analysis. Brain Res Brain Res Rev, 29(2-3), 169–195. 10.1016/s0165-0173(98)00056-3

Klimesch, W., Sauseng, P., & Hanslmayr, S. (2007). EEG alpha oscillations: The inhibition-timing hypothesis. In Brain Research Reviews (Vol. 53, pp. 63–88).

Levit, R. A., Sutton, S., & Zubin, J. (1973). Evoked potential correlates of information processing in psychiatric patients. Psychological Medicine, 3(4), 487–494. 10.1017/S0033291700054295

Liesefeld, H. R., & Muller, H. J. (2019). Distractor handling via dimension weighting. Curr Opin Psychol, 29, 160–167. 10.1016/j.copsyc.2019.03.003

Liston, C., Matalon, S., Hare, T. A., Davidson, M. C., & Casey, B. J. (2006). Anterior cingulate and posterior parietal cortices are sensitive to dissociable forms of conflict in a task-switching paradigm. Neuron, 50(4), 643–653. 10.1016/j.neuron.2006.04.015

Lukas, S., Philipp, A. M., & Koch, I. (2010). Switching attention between modalities: further evidence for visual dominance. Psychol Res, 74(3), 255–267. 10.1007/s00426-009-0246-y

Luks, T. L., Simpson, G. V., Feiwell, R. J., & Miller, W. L. (2002). Evidence for Anterior Cingulate Cortex Involvement in Monitoring Preparatory Attentional Set. NeuroImage, 17(2), 792–802. 10.1006/nimg.2002.1210

Maris, E., & Oostenveld, R. (2007). Nonparametric statistical testing of EEG- and MEG-data. Journal of Neuroscience Methods, 164(1), 177–190. 10.1016/j.jneumeth.2007.03.024

Megevand, P., Molholm, S., Nayak, A., & Foxe, J. J. (2013). Recalibration of the multisensory temporal window of integration results from changing task demands. PLoS ONE, 8(8), e71608. 10.1371/journal.pone.0071608

Miller, J. (1982). Divided attention: evidence for coactivation with redundant signals. Cogn Psychol, 14(2), 247–279. 10.1016/0010-0285(82)90010-x

Molholm, S., Ritter, W., Murray, M. M., Javitt, D. C., Schroeder, C. E., & Foxe, J. J. (2002). Multisensory auditory-visual interactions during early sensory processing in humans: a high-density electrical mapping study. Brain Res Cogn Brain Res, 14(1), 115–128. 10.1016/s0926-6410(02)00066-6

Monsell, S. (2003). Task switching. Trends Cogn Sci, 7(3), 134–140. 10.1016/s1364-6613(03)00028-7

Muller, H. J., Reimann, B., & Krummenacher, J. (2003). Visual search for singleton feature targets across dimensions: Stimulus- and expectancy-driven effects in dimensional weighting. J Exp Psychol Hum Percept Perform, 29(5), 1021–1035. 10.1037/0096-1523.29.5.1021

Orr, J. M., & Weissman, D. H. (2009). Anterior cingulate cortex makes 2 contributions to minimizing distraction. Cereb Cortex, 19(3), 703–711. 10.1093/cercor/bhn119

Pascual-marqui, R. D., Gonzalez-andino, S. L., & Valdes-sosa, P. A. (1988). Current Source Density Estimation and Interpolation Based on the Spherical Harmonic Fourier Expansion. International Journal of Neuroscience, 43(3-4), 237–249. 10.3109/00207458808986175

Perrin, Pernier, Bertrand, & Echallier. (1989). Spherical splines for scalp potential and current density mapping. Electroencephalography and Clinical Neurophysiology, 72, 184–187.

Picton, T. W., Hillyard, S. A., Krausz, H. I., & Galambos, R. (1974). Human auditory evoked potentials. I. Evaluation of components. Electroencephalogr Clin Neurophysiol, 36(2), 179–190. 10.1016/0013-4694(74)90155-2

Plochl, M., Fiebelkorn, I., Kastner, S., & Obleser, J. (2022). Attentional sampling of visual and auditory objects is captured by theta-modulated neural activity. Eur J Neurosci, 55(11-12), 3067–3082. 10.1111/ejn.15514

Poole, D., Miles, E., Gowen, E., & Poliakoff, E. (2021). Shifting attention between modalities: Revisiting the modality-shift effect in autism. Atten Percept Psychophys, 83(6), 2498–2509. 10.3758/s13414-021-02302-4

Posner, M. I., Nissen, M. J., & Klein, R. M. (1976). Visual dominance: An information-processing account of its origins and significance. Psychological Review, 83(2), 157–171. 10.1037/0033-295x.83.2.157

Post, L. J., & Chapman, C. E. (1991). The Effects of Cross-Modal Manipulations of Attention on the Detection of Vibrotactile Stimuli in Humans. Somatosensory & Motor Research, 8(2), 149–157. 10.3109/08990229109144739

Rist, F., & Cohen, R. (1987). Effects of modality shift on event-related potentials and reaction time of chronic schizophrenics. Electroencephalogr Clin Neurophysiol, 40(0424-8155 (Print)), 738–745.

Shaw, L. H., Freedman, E. G., Crosse, M. J., Nicholas, E., Chen, A. M., Braiman, M. S., Molholm, S., & Foxe, J. J. (2020). Operating in a Multisensory Context: Assessing the Interplay Between Multisensory Reaction Time Facilitation and Inter-sensory Task-switching Effects. Neuroscience, 436, 122–135. 10.1016/j.neuroscience.2020.04.013

Shenhav, A., Botvinick, M. M., & Cohen, J. D. (2013). The expected value of control: an integrative theory of anterior cingulate cortex function. Neuron, 79(2), 217–240. 10.1016/j.neuron.2013.07.007

Sohn, M. H., Ursu, S., Anderson, J. R., Stenger, V. A., & Carter, C. S. (2000). The role of prefrontal cortex and posterior parietal cortex in task switching. Proc Natl Acad Sci U S A, 97(24), 13448–13453. 10.1073/pnas.240460497

Spence, C., Nicholls, M. E., & Driver, J. (2001). The cost of expecting events in the wrong sensory modality. Percept Psychophys, 63(2), 330–336. 10.3758/bf03194473

Tollner, T., Gramann, K., Muller, H. J., & Eimer, M. (2009). The anterior N1 component as an index of modality shifting. J Cogn Neurosci, 21(9), 1653–1669. 10.1162/jocn.2009.21108

Turatto, M., Benso, F., Galfano, G., & Umilta, C. (2002). Nonspatial attentional shifts between audition and vision. J Exp Psychol Hum Percept Perform, 28(3), 628–639. 10.1037//0096-1523.28.3.628

Uppal, N., Foxe, J. J., Butler, J. S., Acluche, F., & Molholm, S. (2016). The neural dynamics of somatosensory processing and adaptation across childhood: a high-density electrical mapping study. J Neurophysiol, 115(3), 1605–1619. 10.1152/jn.01059.2015

Vallat, R. (2018). Pingouin: statistics in Python. Journal of Open Source Software, 3(31). 10.21105/joss.01026

Verleger, R., & Cohen, R. (1978). Effects of certainty, modality shift and guess outcome on evoked potentials and reaction times in chronic schizophrenics. Psychological Medicine, 8(1), 81–93. 10.1017/S0033291700006656

Wylie, G., & Allport, A. (2000). Task switching and the measurement of “switch costs”. Psychol Res, 63(3-4), 212–233. 10.1007/s004269900003

Wylie, G. R., Javitt, D. C., & Foxe, J. J. (2003). Task switching: a high-density electrical mapping study. NeuroImage, 20(4), 2322–2342. 10.1016/j.neuroimage.2003.08.010

Wylie, G. R., Javitt, D. C., & Foxe, J. J. (2004). Don’t think of a white bear: an fMRI investigation of the effects of sequential instructional sets on cortical activity in a task-switching paradigm. Hum Brain Mapp, 21(4), 279–297. 10.1002/hbm.20003

Wylie, G. R., Murray, M. M., Javitt, D. C., & Foxe, J. J. (2009). Distinct neurophysiological mechanisms mediate mixing costs and switch costs. J Cogn Neurosci, 21(1), 105–118. 10.1162/jocn.2009.21009

